# MrDNA: A multi-resolution model for predicting the structure and dynamics of nanoscale DNA objects

**DOI:** 10.1101/865733

**Authors:** Christopher Maffeo, Aleksei Aksimentiev

## Abstract

Although the field of structural DNA nanotechnology has been advancing with an astonishing pace, *de novo* design of complex 3D nanostructures remains a laborious and time-consuming process. One reason for that is the need for multiple cycles of experimental characterization to elucidate the effect of design choices on the actual shape and function of the self-assembled objects. Here, we demonstrate a multi-resolution simulation framework, `mrdna`, that, in 30 minutes or less, can produce an atomistic-resolution structure of an arbitrary DNA nanostructure with accuracy on par with that of a cryo-electron microscopy (cryo-EM) reconstruction. We demonstrate fidelity of our `mrdna` framework through direct comparison of the simulation results with the results of cryo-EM reconstruction of multiple 3D DNA origami objects. Furthermore, we show that our approach can characterize an ensemble of conformations adopted by dynamic DNA nanostructures, the equilibrium structure and dynamics of DNA objects constructed using a self-assembly principle other than origami, i.e., wireframe DNA objects, and to study the properties of DNA objects under a variety of environmental conditions, such as applied electric field. Implemented as an open source Python package, our framework can be extended by the community and integrated with DNA design and molecular graphics tools.

## 1 Introduction

DNA self-assembly has emerged as a versatile and robust approach to constructing nanoscale systems ^1–8^. Since its conception^1^, methods for designing and synthesizing self-assembled DNA nanostructures have been progressing steadily^2,9–14^, advancing to two-dimensional origami nanostructures^4^, to three-dimensional origami^5^ and brick^15^ assemblies, to polygon mesh wireframes^16,17^. The ease with which one can customize and functionalize DNA nanoscale objects have catalyzed proof-of-concept developments of a wide range of novel materials^18^ and functional objects, including sensors for pH^19^, voltage^20^ and force^21^; nanoscale containers^22,23^; masks for nanolithography^24,25^; and scaffolds for arranging nanotubes and nanoparticles^26,27^. Presently, gigadalton three-dimensional objects can be constructed through hierarchical self-assembly of DNA molecules^28,29^ with single-nucleotide precision, which makes DNA nanostructures uniquely amenable for mastering spatial organization at the nanoscale^28,29^.

Computational prediction of the *in situ* properties of DNA nanostructures can greatly facilitate the nanostructure design process. Several computational models of DNA nanostructures have been developed already, varying by the amount of detail provided by the computational description. On the coarse end of the spectrum, the CanDo finite element model^30^ minimizes the mechanical stress within a DNA origami structure exerted by its pattern of crossovers to provide a fast estimate of its preferred conformation. At the opposite end of the spectrum, all-atom molecular dynamics (MD) simulations have been used to study the structure^31–33^, and conductance^34–37^ of DNA nanostructures and channels at a much higher computational cost. In between, a number of general purpose coarse-grained (CG) models of DNA are available^38–41^, but only very few have been applied specifically to the study of DNA nanostructures^32,42,43^ including the CG oxDNA^42^ model and all-atom, implicit solvent ENRG-MD^32^ methods.

A major objective for computational models of DNA nanostructures is to give the designers quick feedback on the impact of their design choices. The models employed by the coarsest structure prediction tools may not adequately represent DNA, for example, to guide the placement of fluorescent dyes or functionalized groups. Coarse-grained models do not so far provide insights into the ion conducting and rheological properties of DNA nanostructures. To date, only all-atom MD has been used to computationally examine these properties. On the other hand, higher resolution modeling can be prohibitively slow for routine structural characterization, or for sampling the configurational space of a nanostructure, particularly when the equilibrium, relaxed conformation of an object departs strongly from its initial idealized configuration as obtained from popular CAD tools such as cadnano^44^, vHelix^16^ or DAEDALUS^17^.

Here we present a computational framework that combines low- and high-resolution models of DNA objects to enable fast and accurate predictions of the objects’ structure and dynamics suitable for iterative design applications, Fig. 1. Starting from an idealized design of a DNA object, our structure prediction protocol performs rapid relaxation of the design using a low-resolution model. The outcome of the relaxation simulation is further refined through several simulations performed at increasing resolution to produce an accurate, fully atomistic representation of the object’s *in situ* structure. Our computational framework can also characterize the conformational dynamics of the object at multiple time scales and characterize the behavior of the objects in the presence of external potentials and/or under non-equilibrium conditions.

**Figure 1:**
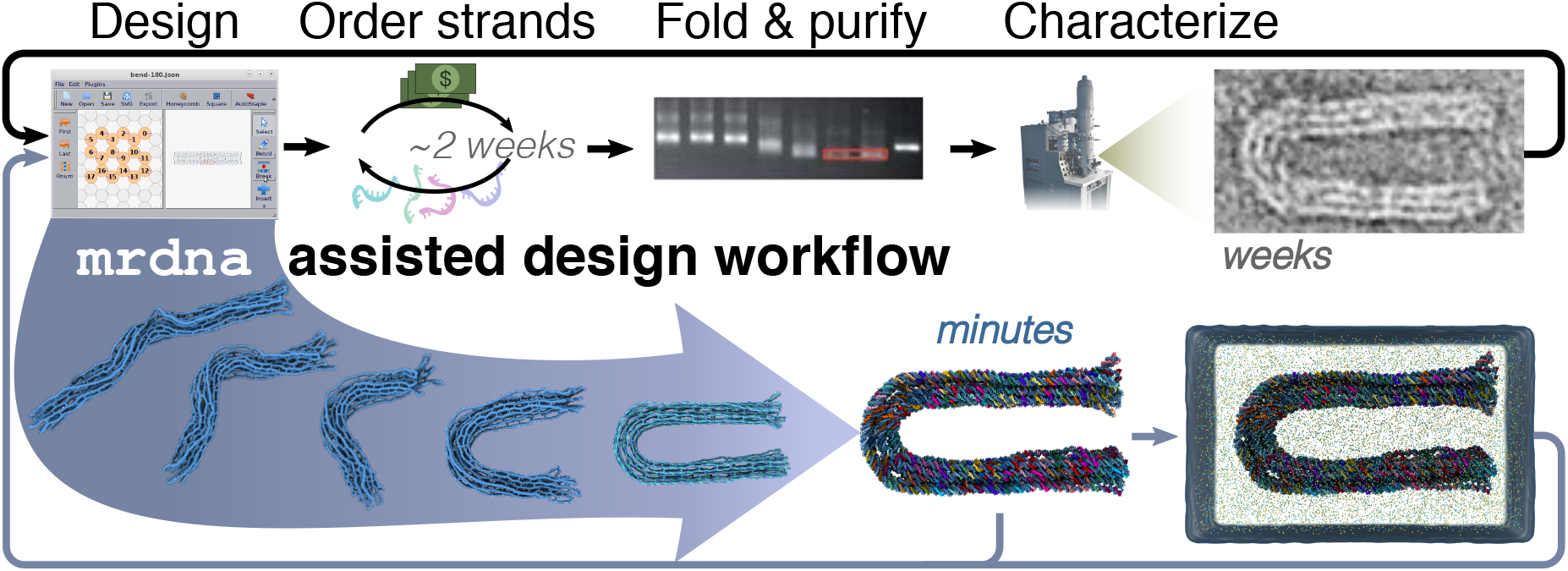
DNA nanostructure design workflow augmented by the multi-resolution modeling framework, `mrdna`. The top row depicts a traditional design cycle involving several steps that result in a latency of several weeks before feedback is obtained. A DNA origami bundle with a programmed 180° bend^45^ is shown as an example. The bottom row illustrates how `mrdna` can be used to obtain the equilibrium *in situ* configuration of the DNA nanostructure. The snapshots within the arrow illustrate several stages of a CG simulation trajectory; the structure immediately to the right of the arrow shows the final all-atom configuration. The resulting all-atom structures are directly suitable for subsequent all-atom MD simulations, the right most image shows an example of a solvated all-atom system. TEM images reproduced from Ref. 45 with permission.

## 2 MATERIALS AND METHODS

All CG simulations of DNA origami objects were performed using an in-house developed GPU-accelerated CG simulation package, Atomic Resolution Brownian Dynamics (ARBD)^46^. ARBD supports Brownian and Langevin dynamics simulations of systems composed of isotropic point-like particles interacting through tabulated bonded and non-bonded interactions. ARBD simulations are currently configured through system-specific files that the `mrdna` framework writes automatically.

### 2.1 Overview of the mrdna framework

The `mrdna` framework is shipped as a Python package that describes DNA nanostructures using several levels of abstraction. At the base level are a collection of classes that represent collections of beads or atoms interacting through user-specified bonded and non-bonded interactions. These classes provide a file-based interface to our simulation engine, ARBD. The second level of abstraction provides classes that describe contiguous double-stranded and single-stranded regions or “segments” of DNA that can be joined together through intrahelical, terminal and crossover connections. The spatial configuration of each DNA segment is described by a spline function defined in cartesian coordinates. For dsDNA, a four-dimensional spline fit through quaternions optionally provides a parametric representation of the local orientation throughout the duplex. The use of splines allows the classes to generate bead-based models with user-specified resolution. Alternatively the segment classes can generate an atomistic or oxDNA model of the structure. The spline representation mediates conversion from one resolution to another; after a CG simulation is performed, new splines are generated that pass through the coordinates of the beads at the end of the simulation.

The segment classes can be used directly for algorithmic design of DNA nanostructures or can be used to write scripts that automatically convert a model from another software package into an `mrdna` model. For example, a class was developed using the new cadnano2.5 Python API to read in a cadnano json design file, allowing direct access to the cadnano data structures for conversion to the segment-based model. Similar classes have been developed for converting vHelix mesh models and atomic PDBs into `mrdna` models.

### 2.2 Conversion of spline model to bead model

Beads representing interaction sites within double- and single-stranded segments are iteratively placed through the structure and assigned a position within the segment. First, beads are placed at the middle of every intrahelical (or intrastrand) connection. Beads are next placed on both sides of each crossover, When two beads are intrahelically adjacent and placed within the user-provided resolution of the model, they are merged into one bead. Next, beads are placed at the ends of each segment if none already exists. Beads are placed between each pair of existing beads so that the density of beads is lower than a user-specified threshold. Finally, the intrahelical distance between adjacent beads is calculated from their positions and used to assign a nucleotide count to each bead. The beads are hierarchically clustered according to nucleotide count, ensuring a tractable number of bead types is provided to the simulation engine.

### 2.3 Variable-resolution bead model of DNA nanostructures

Once beads have been placed throughout a DNA nanostructure, bonded and non-bonded potentials are specified to describe their interactions. A tabulated nonbonded potential was developed by iteratively tuning interactions until the osmotic pressure in an array of 256 two-turn DNA helices— each made effectively infinite through periodic boundary conditions—matched the experimental measurements of Rau and Parsegian for DNA in a 25 mM MgCl_2_ electrolyte^47^, see Fig. S1. If the user supplies a custom Debye length, a correction is applied to the tabulated nonbonded potential corresponding to the difference of Yukawa potentials for the user-supplied and 11.1 Å Debye lengths, the latter representing the ion condition of the Rau and Parsegian data. The nonbonded interaction between any pair of beads is scaled by the nucleotide count of each bead involved. DNA array simulations verified that the pressure was insensitive to the resolution of DNA under 1, 3, and 5 bp/bead conditions, with and without a local representation of the twist. The same potential is used for nonbonded interactions involving ssDNA. For dsDNA, nonbonded exclusions are placed between all pairs of beads intrahelically within 20-bp of one another. For ssDNA, exclusions are placed between pairs of beads in the same strand that are within five nucleotides (nts).

The bonded interactions within the model are realized by harmonic potentials with spring constants and rest lengths determined from known polymer properties of DNA, Fig. S2. A harmonic bond connects each consecutive pair of beads in a helix or strand with a separation-dependent rest length (3.4 Å/bp or 5 Å/nt) and a spring constant derived from the elastic moduli of dsDNA and ssDNA (1000 and 800 pN, respectively)^48^. A harmonic angle potential is placed on the angle formed by the bonds between consecutive pairs of beads within a helix or strand with a 0° rest angle and spring constant *k_p_* derived from the persistence length of DNA. Specific values of *k_p_* were obtained by numerically solving the following equation:

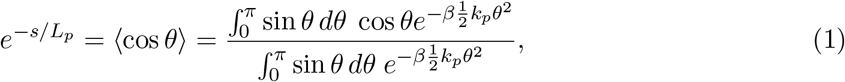

where *s* is a half of the contour length between the first and the third bead, *L_p_* is the persistence length, *θ* is the angle between the beads, *k_p_* is the spring constant used in the simulation, and *β* = 1/*k_B_T*, where *k_B_T* is the thermodynamic temperature.

The `mrdna` models can be generated with or without a local representation of the orientation of each basepair. When the twist is represented locally, an orientation bead representing the major groove is placed alongside each dsDNA backbone bead, attached to its parent bead by a harmonic potential (1.5 Å rest length; *k*_spring_ = 30 kcal mol^−1^ Å^−2^). A harmonic angle potential (90° rest angle; *k*_spring_ = 0.5 *k_p_*) is placed between consecutive intrahelical beads and each of the orientation beads. A harmonic potential is placed on the dihedral angle formed by each orientation bead, its parent intrahelical bead, the adjacent intrahelical bead and the corresponding orientation bead with a separation-dependent rest angle (10.44 helical rise) and a spring constant *k*_tw_(*s*) designed to produce a 90-nm twist-persistence length for dsDNA to match the measured twist persistence length under tension^49^, see Figs. S2, S3.

Finally, a harmonic bond is placed between intrahelical beads on either side of a junction spanning two helices (such as a DNA origami crossover connection). The spring constant (4 kcal mol^−1^ Å^−2^) and the rest length (18.5 Å) of the bond were derived from the distribution of crossover distances observed in atomistic simulations^33^. When neither side of the junction occurs at the end of a DNA helix, the junction is assumed to represent a DNA origami crossover, and a heuristic dihedral angle potential is placed on an intrahelical bead on one side of the first junction bead, the first junction bead, the second junction bead, and the intrahelical bead on the opposite side of the junction (*k*_spring_ = 0.25 *k_p_*; rest angle 180°) to keep the adjacent helices roughly parallel. If twist is locally represented, an additional harmonic potential is added to the dihedral angle formed on each helix by the junction bead on the opposite helix, the junction bead, the bead adjacent to the junction bead, and the junction bead’s orientation bead (*k*_spring_ = 0.25; *k_p_*; rest angle = ±120, depending on the strand) to ensure that the connecting strand on each side of the junction faces the helix on the other side of the junction, Fig. S4A. In the event that twist is not locally represented, the torsion within each helix is described by a dihedral angle potential applied between the beads forming consecutive junctions in the helix. The spring constant and rest angle are calculated as described above for the local representation of twist, except if the junctions occur on different strands, 120° will be added to (removed from) the rest angle if the 5’-to-3’ direction at each junction site points away from (towards) the other junction, Fig. S4B.

## 3 RESULTS

### 3.1 Multi-resolution model of self-assembled DNA nanostructures

The `mrdna` framework performs an automatic multi-stage simulation of a DNA nanostructure starting from its idealized initial configuration, which can come from a variety of sources. Here, we illustrate the basic workflow of `mrdna` using, as an example, a cadnano^44^ design of a curved six-helix bundle.

Starting from a configuration file generated by the design software, Fig. 2A, the standard `mrdna` relaxation procedure begins with the construction of a low-resolution, 5 basepair (bp)/bead CG model, Fig. 2B. The nonbonded interactions between the beads are described by empirical potentials that have been calibrated to reproduce the experimentally measured osmotic pressure of a DNA array in 25 mM MgCl_2_ electrolyte^47^, Fig. S1. Our model can accommodate other ionic strength conditions at the level of the Debye-Hückel approximation, accounting for ionic strength conditions through a Debye length parameter. The beads within each double- or single-stranded region are connected by harmonic bond and angle potentials with separation-dependent rest-lengths and spring constants derived to match the experimentally measured elastic moduli and persistence lengths, Figs. S2, S3. Finally, bond, angle and dihedral potentials were defined for the beads at each junction, see Methods for further details. The resulting model is relaxed from its initial, idealized geometry using the simulation engine, ARBD^46^. Figure 2B-D illustrate the relaxation simulation of the curved six-helix bundle.

**Figure 2:**
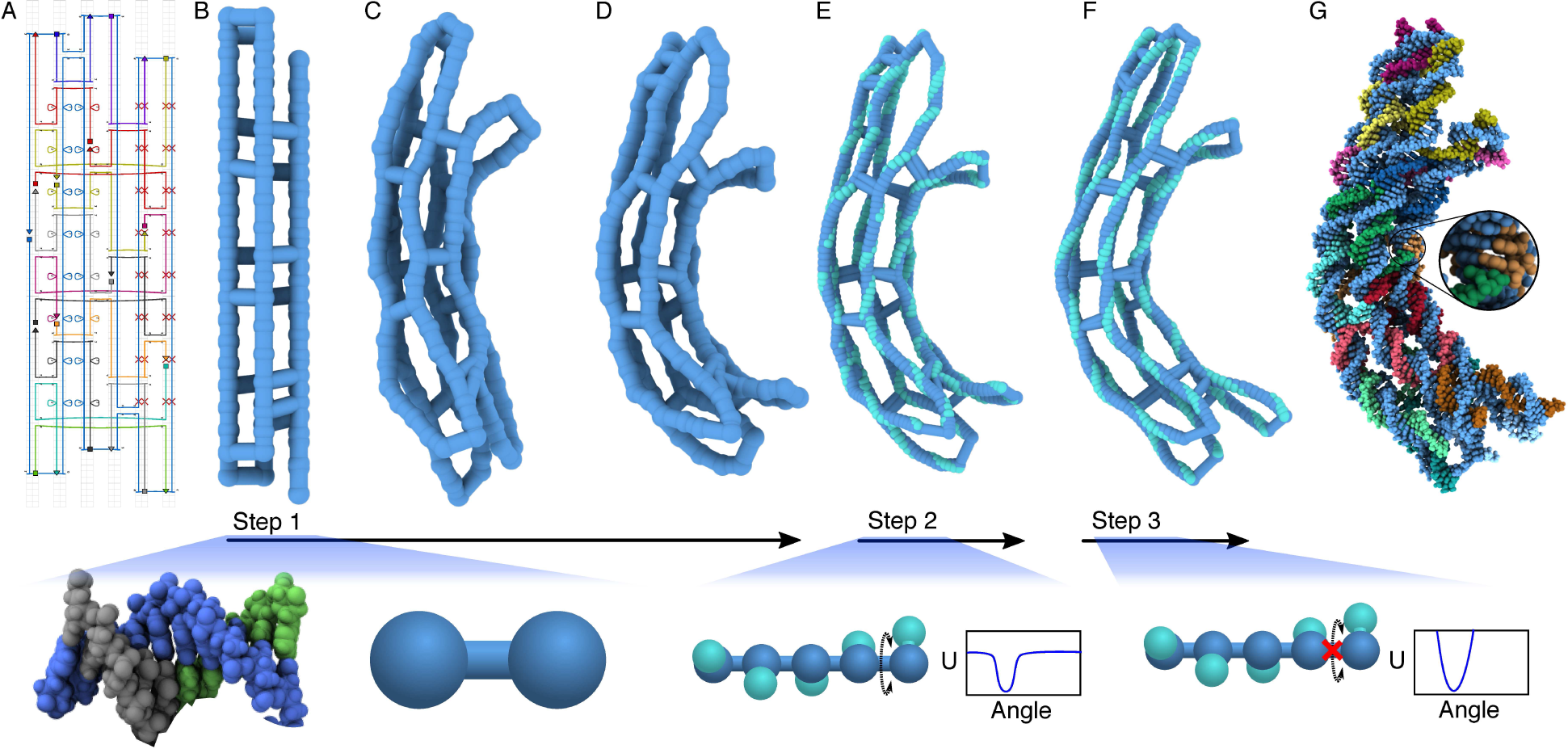
Multiresolution structural relaxation of a curved DNA origami object. (A) Cadnano^44^ design of a curved six-helix bundle. A regular pattern of insertions in two of the helices and deletions in two opposing helices induce the curvature^45^. (B-D) Low-resolution CG simulation of the bundle. In just 40 nanoseconds, the bundle adopts a curved conformation. The bundle is shown using a bead-and-stick representation, where each bead represents five basepairs, on average. (E,F) High-resolution CG simulations of the bundle. A spline-based mapping procedure yields a high-resolution (2-bead per basepair) model by interpolation. The high-resolution model includes a local representation of the each basepair’s orientation (teal beads). The twist dihedral angle potential between adjacent orientation beads is smoothly truncated to allow the linking number to relax during a brief, 10-ns simulation (E). A subsequent 10-ns simulation performed without the truncation of the twist potential allows complete relaxation of the bundle (F). (G) Resulting atomic model of the curved six-helix bundle. Canonical basepairs are placed throughout the structure using the spline-based mapping procedure. The images below each of the three CG simulation steps illustrate schematically how the model represents one turn of DNA.

The structure obtained at the end of the above relaxation process is further refined through additional simulations performed using models of increasing resolution. To facilitate the change to a higher (2 bead/bp) resolution, a spline function is first defined to trace the center line of each double- or single-stranded DNA region in the structure obtained at the end of the low-resolution simulation, Fig. 2D. The spline function is used to find the coordinates of the finer resolution model by interpolation, Fig. 2E. Two beads are used to represent each basepair of dsDNA: one bead representing the basepair’s center of mass and one represent the basepair’s orientation—the location of the major groove. The initial placement of the orientation beads is equilibrated in a brief simulation using a harmonic dihedral potential that is smoothly truncated at 1 kcal mol^−1^, which allows the linking number of DNA to change during this equilibration simulation. Following that, the model is simulated using a full (non-truncated) harmonic twist potential, producing a high-resolution equilibrated structure, Fig. 2F. Center line and orientation spline fits through the average coordinates obtained at the end of the 2 beads/bp equilibration simulation are used to construct an all-atom structure, Fig. 2G. The Methods section provides a complete description of the mapping and model parameterization procedures.

### 3.2 Structure prediction

Using the `mrdna` framework, one can obtain an equilibrium, fully atomistic structure of a complex DNA origami object with the accuracy of a cryo-electron microscopy (cryo-EM) reconstruction in 30 minutes or less. To demonstrate the accuracy of the model, the “pointer” DNA origami object^50^ was simulated for 400 ns (2 million steps) at 5-bp/bead resolution, requiring ~30 min on an NVIDIA RTX 2080 GPU, see Supporting Animation 1. The resulting conformations were compared with the high-resolution three-dimensional reconstruction obtained using cryo-EM^50^.

To enable comparison of the structures, the simulated configurations were first aligned to the initial, idealized configuration by minimizing the root mean squared deviations (RMSD) of the beads’ coordinates. The bead coordinates from the final 200-ns fragment of the simulation were then averaged, producing a simulation-derived structure, shown in blue in Fig. 3A. For visual comparison, the resulting time-averaged 2-bead/bp structure was docked into the cryo-EM density (white) using the brute-force voltool fit algorithm of VMD^51^, revealing a good overall match for the global configuration, Fig. 3A (top row). To quantitatively monitor the relaxation of the structure in the `mrdna` simulation, we used, as a reference, the all-atom model^50^ produced by flexible fitting of the idealized conformation into the cryo-EM density^52^. The CG trajectory was converted to an all-atom representation using our multiresolution simulation procedure, where two very brief (0.5 ns) simulations at 2-bead/bp resolution were run prior to all-atom conversion, which largely preserved the instantaneous configurations sampled by the 5-bp/bead simulation. In the first of the mapping simulations, the linking number of the DNA was allowed to relax.

**Figure 3:**
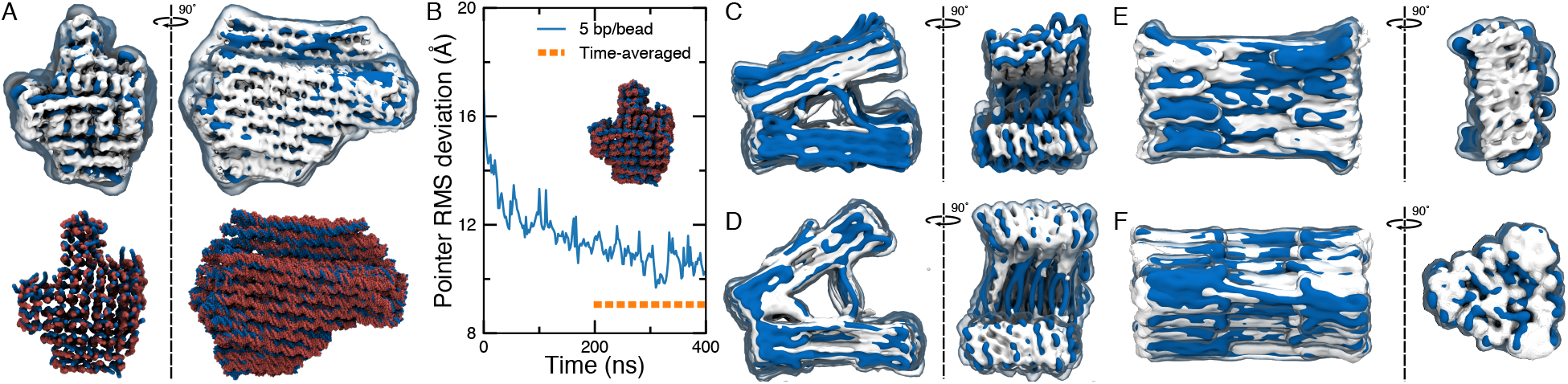
Structure prediction with `mrdna`. (A) Comparison of the structural models obtained through an `mrdna` simulation and cryo-EM reconstruction (EMD-2210) of the “pointer” structure^50^. The top row depicts the simulated and experimental structures as isosurfaces of the CG bead (blue, 1 bp/Å^3^ isovalue) and electron (white, 0.08 isovalue) densities, respectively. The bottom row compares the simulated (blue) and experimental (red) all-atom models. In the top row, simulated structural fluctuations are visualized using a semi-transparent surface showing the average density of CG beads during the final 200 ns of a 5-bp/bead simulation. Note that some helices along the periphery of the structure were not resolved in the cryo-EM model^50^. (B) Root-mean-square deviation (RMSD) of the C1′ atom coordinates of the scaffold strand from those of the cryo-EM-derived atomic structure^50^ during `mrdna` relaxation of the pointer structure at 5-bp/bead resolution. For RMSD calculations, instantaneous configurations were mapped to an all-atom model. The orange dashed line shows the RMSD between the cryo-EM structure and the atomic structure time-averaged over the final 200-ns. (C-F) Comparison of the `mrdna` (blue, 1 bp/Å^3^ isovalue) and cryo-EM (white, 0.02 isovalue) models of the objects designed and characterized by the Dietz group^28^: v-brick structures without (C; EMD-3828) and with (D; EMD-3828) twist correction, the connector block (E; EMD-3827) and the triangular vertex (F; EMD-3826).

The RMSD of the scaffold C1′ atoms from the experimentally derived structure was seen to decrease rapidly at first and more steadily as the simulation progressed, Fig. 3B. After 200 ns, the RMSD was decreasing rather slowly, but still exhibited significant fluctuations. Such fluctuations of the RMSD value reflect the equilibrium fluctuations of the underlying structure, which are averaged out in the cryo-EM reconstruction. To remove the effect of the fluctuations, we aligned and averaged the all-atom structures over the last 200 ns of the `mrdna` simulation. The resulting all-atom structure matched the cryo-EM derived all-atom model very closely, bottom row of Fig. 3A, and the RMSD between the simulated and experimental structures dropped below 1 nm, dashed line in Fig. 3B. Thus, the `mrdna` simulation resulted in a model with comparable deviation from the experimentally-derived structure (9.06 Å) as the more detailed models, ENRG-MD^32^ (9.12 Å) and oxDNA^39,40^ (9.41 Å) but at a fraction of the computational cost. The oxDNA comparison was obtained by converting the second half of a billion-step oxDNA trajectory of the pointer structure that we performed to all-atom coordinates using tacoxDNA^53^ and averaging over the ensemble of the all-atom structures. It it worth noting that all of the above models yielded exceptional agreement with experiment, considering the reported 1.15-nm resolution of the reconstruction.

Further validation of the `mrdna` structure prediction protocol was obtained by simulating a set of four honeycomb lattice structures that were recently designed and characterized through 3D cryo-EM reconstructions by the Dietz group^28^. The structures included “v-brick” objects, without and with twist correction, a rectangular prism “connector”, and a triangular prism “vertex”. The protocol described above for the pointer structure was applied to each object, resulting in average models that could be docked very nicely in the corresponding cryo-EM densities, Fig. 3 C-F. The ends of the bricks, excepting the vertex structure, were seen to spread slightly more in `mrdna` simulation than in the cryo-EM reconstructions. However, the overall simulated shape—including twist within each prism and, where applicable, the skew between connected prisms—matched the cryo-EM density very closely, Supporting Animation 2-6.

Many DNA origami objects are designed to bend or twist via a pattern of basepair insertions and deletions^45^. Atomistic (or ENRG-MD) simulations of such a structure that start from the idealized configuration can become trapped in a local energy minimum, preventing complete relaxation of the object. The `mrdna` framework was used to relax an ensemble of curved and twisted structures, Fig. 4 B,C. All of the objects relaxed to configurations that closely resemble the experimental transmission electron microscopy (TEM) images, providing qualitative validation of the model. However, the object with the 180° bend was deformed slightly out of plane.

**Figure 4:**
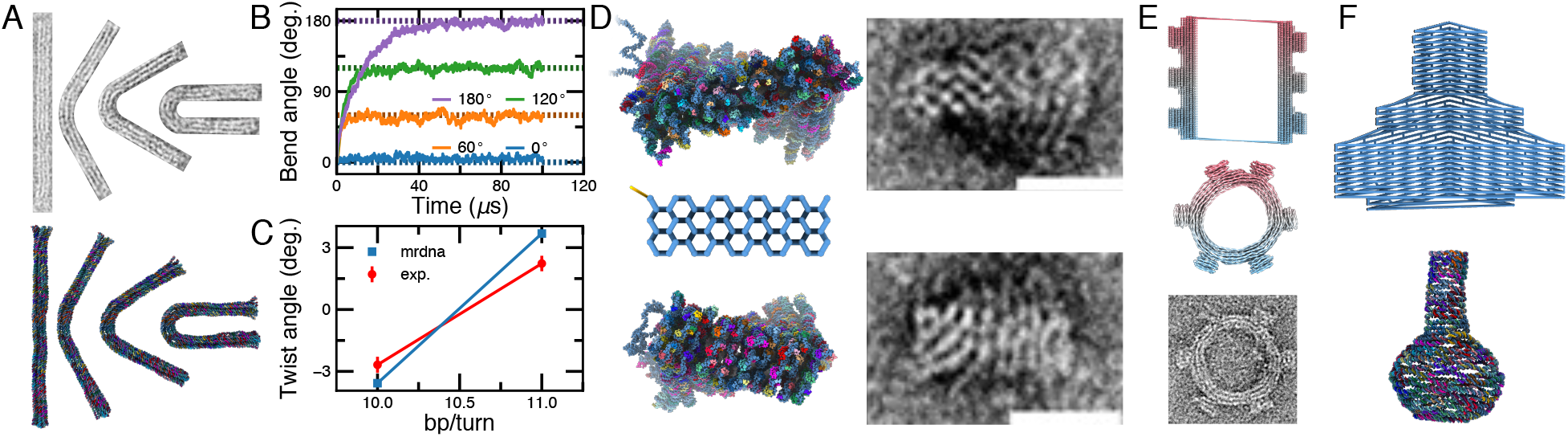
Structural models of curved and twisted DNA origami nanostructures. (A) Structures with programmed bends as characterized by TEM in Ref. 45 (top row) and reconstructed at atomic resolution using `mrdna` (bottom row). From left-to-right, the structures were designed to have a bend of 0°, 60°, 120° and 180°. (B) Bend angle timeseries during 5-bp/bead `mrdna` relaxation of the structures shown in panel A starting from an initial configuration of a straight rod. Dashed lines depict the target angles. (C) Dependence of the twist angle on the helical rise imposed by crossovers for honeycomb brick origami structures^45^ in 5-bp/bead `mrdna` simulations and as measured using TEM^45^. (D) All-atom models of brick origami structures (left, top and bottom) resulting from `mrdna` simulations starting from an idealized configuration (left, center). The right column shows the corresponding TEM images. (E) Gear nanostructure^45^ before (top; 5 bp/bead) and after (center; 2 bead/bp) a `mrdna` simulation. The gear is depicted using a bead-and-stick representation. The bottom panel depicts the corresponding TEM image. (F) Flask nanostructure^54^ before (top; 5 bp/bead) and after (center; atomistic) an `mrdna` simulation. The top panel shows a bead-and-stick representation including long bonds that span the structure. All TEM images were reproduced from Ref. 45 with permission.

The idealized conformation of many DNA nanostructures can sometimes deviate so dramatically from the object’s equilibrated configuration that CG methods, including `mrdna`, are unable to relax the structure. For example, the flask^54^ object from the Yan group is a highly symmetric, flat DNA object that is quickly crushed by intrahelical connections that span the object, see Supporting Animation 7. To make it possible to model such complicated structures, the `mrdna` framework has been implemented as a Python package, which allows one to script modifications to a model. For example, many designs are intended to self-assemble into larger assemblies. The scripting interface was used to combine two models of the half-gear design^45^, applying a 180° rotation to one of the models and adding intrahelical connections between the two half-gears, Fig. 4 D. Returning to the example of the Yan group’s flask, a second Python script was used to apply a 90° rotation to half of each helix in a DNA flask before running the multiresolution simulations, Fig. 4 E and Supporting Animation 8. In both cases, scripting was necessary to prevent the rapid collapse of bonds that span the initially straight structure, which would result in a crushed, tangled model of the structure. The resulting relaxed structures were seen to resemble TEM images of the objects, Fig. 4D,E.

### 3.3 Conformational dynamics

DNA origami nanostructures are sometimes designed to adopt multiple conformations. The `mrdna` framework can be used to predict the average structure and the conformational fluctuations of such objects. For example, the Dietz and Castro groups have designed nanoscale calipers using DNA origami that allow measurement of intermolecular forces between proteins attached to each arm of the caliper^55, 56^. TEM can be used to determine the distribution of angles of the caliper with and without attached proteins, allowing inference of intermolecular forces. To demonstrate the utility of the `mrdna` framework for sampling conformational space, a set of 500-*μ*s simulations using the caliper design from the Dietz group was performed using a 5-bp/bead model, Fig. 5A and Supporting Animation 9. The angle between the two arms was extracted at each frame of the simulation trajectory, providing the distribution of the angles between the two arms, Fig. 5B. Substantial overlap between the `mrdna`-generated and experimentally-measured distributions was observed, though the `mrdna` distribution is skewed towards more acute angles, likely because of incomplete parameterization of the ssDNA part of the design.

**Figure 5:**
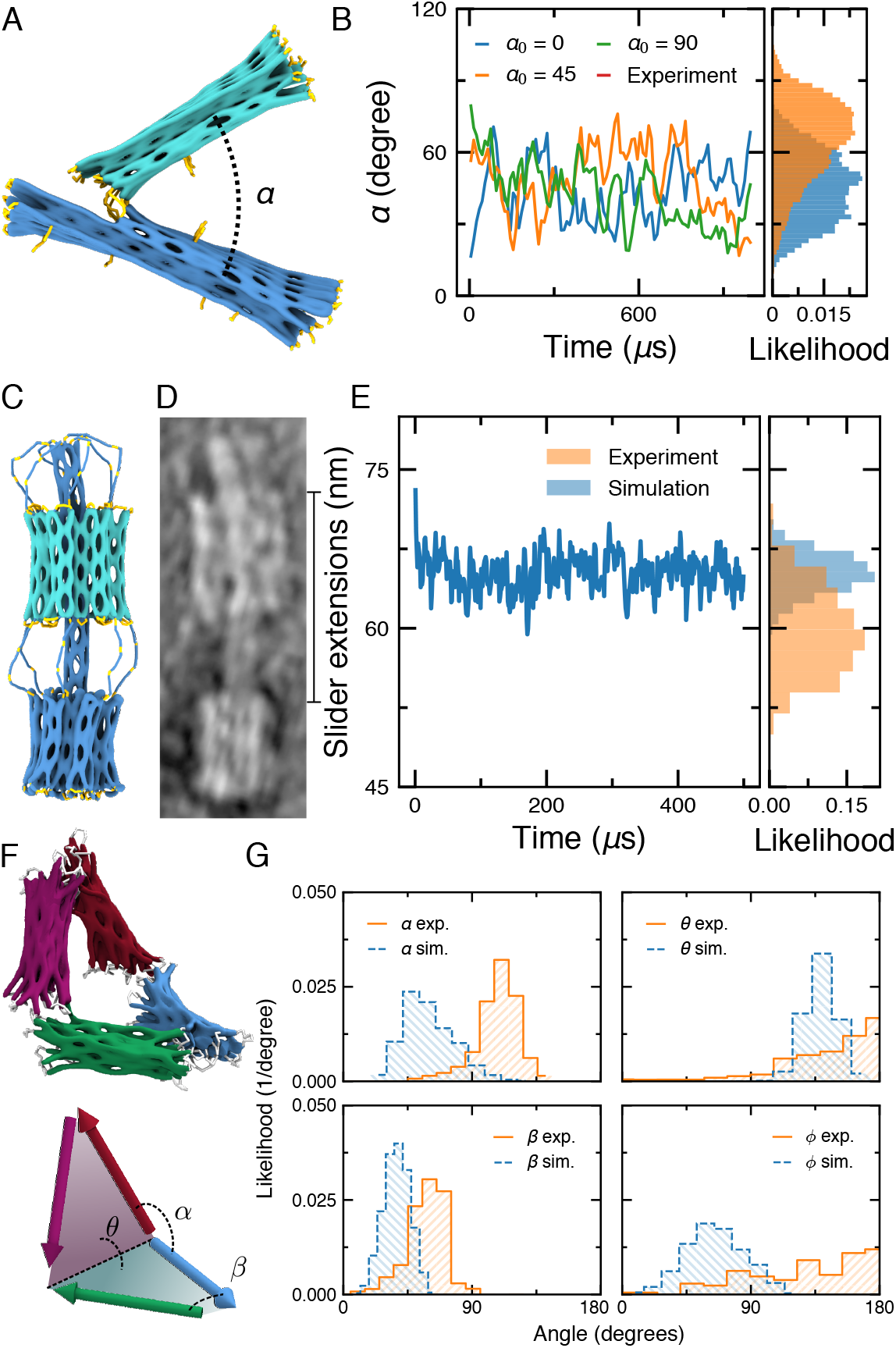
`Mrdna` simulation of dynamic DNA origami nanostructures. (A) Molecular graphics representation of a DNA origami caliper designed by the Dietz group for measuring inter-nucleosome forces ^55^. (B) The angle α between the two arms of the caliper in three 5-bp/bead `mrdna` simulations, starting with the initial angle of 0, 45 or 90 degrees. The caliper angle was computed as the angle between the axes of the arms of the caliper as determined by single-value decomposition^55^. Histograms show the distribution of the simulated angles alongside the distribution extracted from the TEM images^55^. (C,D) Molecular graphics representation (C) and TEM image (D) of a DNA slider nanostructure designed and characterized by the Castro group^57^, reproduced with permission from Ref. 57. (E) The distance between the bearing and the base of the nanostructure during a 5-bp/bead `mrdna` simulation. Histograms show the distribution of the simulated distances alongside the distribution extracted from the TEM images^57^. (F,G) Angle distributions of a Bennett linkage nanostructure designed by the Castro group^58^ in 5-bp/bead `mrdna` simulations. The distributions observed in simulations are plotted alongside the corresponding distributions extracted from single-particle reconstructions of the linkages^59^.

The slider origami nanostructure designed by the Castro group^57^ was simulated next, Fig. 5C,D and Supporting Animation 10. The slider consists of a large base affixed to a six-helix bundle shaft that threads through a movable bearing, which is tethered on opposite ends to the base and to the tip of the shaft by a total of twelve flexible linkers. While the bearing was seen to dwell ~7 nm further from the base structure in `mrdna` simulations with a more narrow distribution than observed experimentally, the position of the bearing is overall in qualitative agreement, Fig. 5E. Interestingly, a similar discrepancy was observed for a similar slider design when it was simulated for a very long time using the oxDNA model^60^. The simulations using the `mrdna` framework could be performed quickly and lasted for about two days.

Finally, we performed simulations of the DNA origami Bennett linkage designed by the Castro group^58^ and imaged using electron tomography^59^, Fig. 5F and Supporting Animation 11. Using the 5-bp/bead model, we were able to sample the full range of motion of the Bennett linkage, revealing stochastic transitions between planar and compact configurations. Distributions for the large and small angles (*α* and *β*, respectively) between adjacent arms and for the large and small dihedral angles (*θ* and *ϕ*, respectively) between the planes formed by adjacent arms were calculated following the overall approach described by the Ren group in a recent study of the object using three-dimensional single-particle reconstructions. Briefly, at each frame of the CG simulation trajectory, the axis of each arm was determined using single-value decomposition. The angles between the arm axes directly provided *α* and *β*. Cross products between the axes of adjacent arms provided normal vectors for the plane formed by the two arms, allowing the dihedral angles *θ* and *ϕ* to be computed from the angle between the normal vectors. The resulting distributions are in approximate agreement with distributions experimentally obtained by the Ren group, Fig. 5G. Hence, our simulation method can be used to quickly study the structural fluctuations of dynamic DNA nanostructures, providing rapid feedback about the mechanics of a design in place of challenging experimental characterization.

### 3.4 Support for non-origami DNA nanostructures

The `mrdna` framework can be used to simulate self-assembled DNA nanostructures produced by approaches other than DNA origami. The same set of commands that are used to construct the spline/bead-based models of cadnano designs can introduce new types of connections between DNA regions, extending the application of the `mrdna` framework to other kinds of DNA nanostructures. For example, the `mrdna` package already includes a code that constructs a multi-resolution model from lists of basepairs, dinucleotide stacks and backbone connections for all the nucleotides in a system. We used that code to simulate a variety of wireframe DNA nanostructures^16^ starting directly from their vHelix Maya designs. The resulting CG MD trajectories characterized the amount of structural fluctuations in the designs and produced representative ensembles of equilibrated all-atom structures, Fig. 6A-E. The results of our `mrdna` simulations of the vHelix objects showed that the overall geometry of each structure was preserved despite relaxation of the DNA.

**Figure 6:**
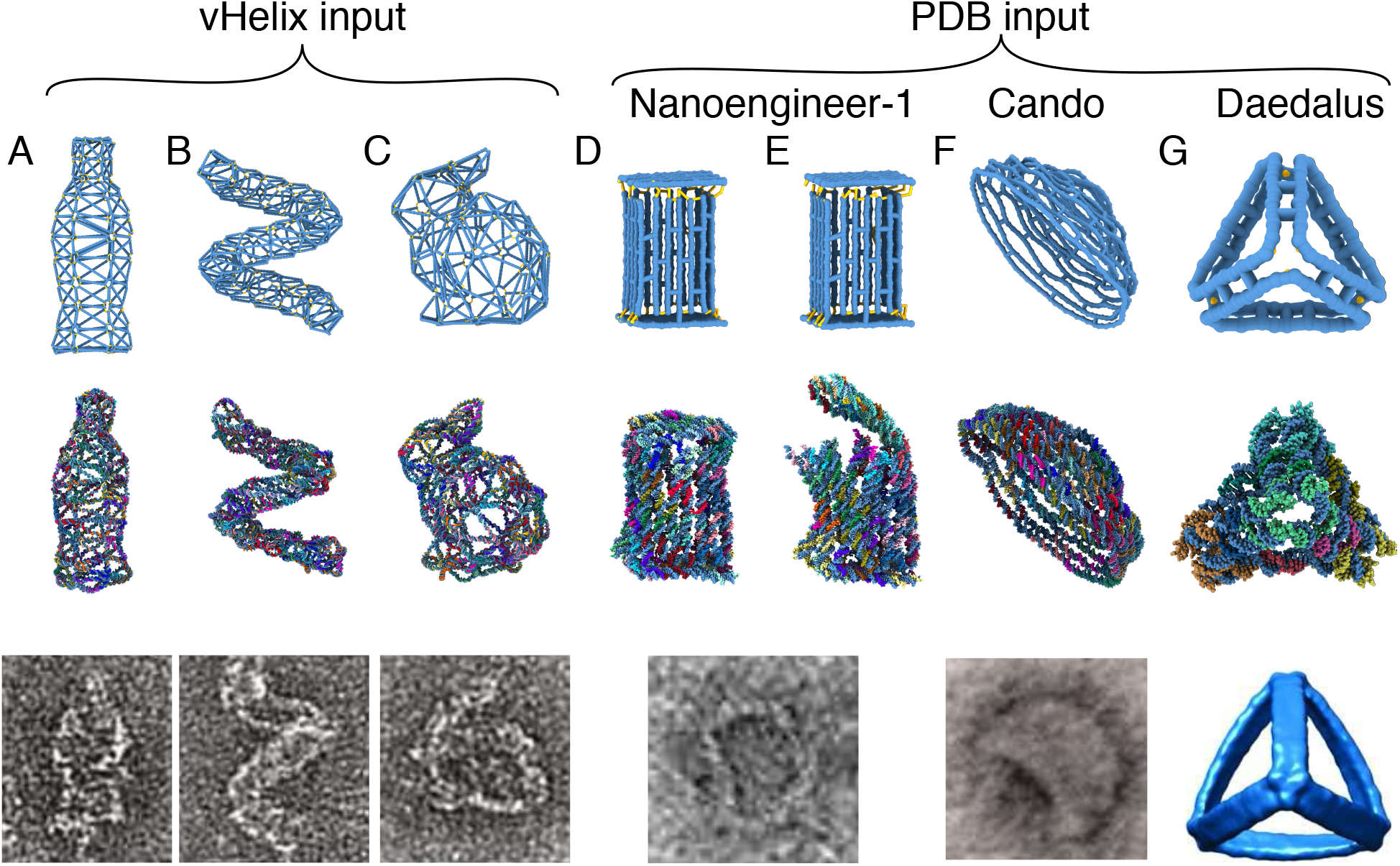
`Mrdna` simulations of DNA nanostructures designed using programs other than cadnano. (A-C) Structural relaxation of vHelix nanostructures starting from the vHelix maya files obtained from the Högberg group^16^. From top-to-bottom, the panels show the configurations of nanostructures prior to the low-resolution CG simulation, the resulting atomistic model, and an experimentally obtained TEM image, reproduced with permission from Ref. 16. (D-G) Structural relaxation of nanostructures using all-atom PDB models as input for `mrdna`. The all-atom PDBs of a box^23^ (D,E), hemisphere^54^ (F) and tetrahedron^17^ (G) were obtained from Nanoengineer-1^61^, CanDo^30,54^ (original design from cadnano), and DAEDALUS^17^. From top-to-bottom, the panels show the configurations of nanostructures prior to the low-resolution CG simulation, the resulting atomistic model, and an experimentally obtained TEM image or reconstruction. The box TEM image is reproduced with permission from Ref. 23. Copyright 2012 American Chemical Society. The hemisphere TEM image is reproduced from Supplementary Figure S1 associated with Ref. 54 with permission from AAAS. The tetrahedron reconstruction is reproduced from Ref. 17 with permission from AAAS.

The `mrdna` framework also includes an algorithm to automatically identify all backbone connections, basepairs, and dinucleotide stacks in an all-atom representation of a DNA nanostructure formatted as a Protein Data Bank (PDB) file. This PDB reader module can import the output of many tools that export PDBs, including CanDo^30^, Nucleic Acid Builder^62^, oxDNA^39,40^ (via the convert_to_atomic.py script distributed with oxDNA), Nanoengineer-1^61^ (via our nanoengi-neer2pdb web service^63^), DAEDALUS^17^, and Tiamat^64^, to automatically generate a corresponding `mrdna` model. We demonstrate this capability here by relaxing several DNA nanostructures using all-atom PDBs from various sources as inputs, Fig. 6F-J. The `mrdna` simulations demonstrated substantial relaxation of the designs away from their idealized initial geometries, with the corresponding final atomistic models having an average RMSD of 24 Å compared to the initial structures. The ability of the `mrdna` framework to read PDB files immediately allows interfacing with a wide array of design and simulation tools, but not all PDB structures are interpreted perfectly by the framework because basepairs or stacks may be assigned incorrectly. Hence, users are cautioned to carefully examine their resulting models to ensure proper conversion.

The atomic configuration resulting from the `mrdna` simulations could be used to inform design modifications or to perform further refinement of the structure using elastic network-guided MD (ENRG-MD)^32^ implicit solvent simulations or explicit solvent all-atom simulations. Moreover, the `mrdna` framework can generate all files needed to perform additional relaxation using the oxDNA model.

### 3.5 Integration with continuum models

DNA nanostructures are usually designed to operate in the environment of a biological system or in the context of a nanotechnological device. In either case, it is useful to be able to couple a model of a DNA nanostructure to other models because a DNA object’s conformation and function can both be influenced by its environment.

The `mrdna` framework uses ARBD^46^, a GPU-accelerated biomolecular simulation engine, which supports mixed models of point particles and rigid body objects as well as three-dimensional grid potentials. Hence, configuration files written by the `mrdna` framework can be manually edited to include interactions with additional entities (e.g. cellular proteins or small molecules) that are not described by the package. The scripting interface for the `mrdna` framework already exposes commands for applying grid-based potentials to DNA nanostructures, allowing one to exert effective forces extracted from continuum models to, for example, simulate DNA nanostructures under the influence of spatially varying electric or plasmonic optical fields^65^.

We demonstrate this capability here by simulating the capture of a DNA nanostructure in a nanopipette, Fig. 7 and Supporting Animation 12. The spatial distribution of the electrostatic potential near the pipette under a 300-mV applied bias was obtained using the Comsol continuum model, as described previously^20^. The potential was applied to each bead in the system with a weight proportional to the number of nucleotides represented by the bead. In our `mrdna` simulation, we observe one leg of the DNA nanostructure to be captured initially, followed by a collapse of the wireframe mesh, which allowed the object to pass fully through the aperture of the pipette. While the above simulation is just an illustrative example, we have previously used similar multiscale/multiphysics simulations to obtain a quantitatively correct description of the voltage-dependent deformation of a DNA nanostructure subject to applied electric field of various magnitudes^20^. We expect the combination of particle-based simulations with continuum modeling to bring much anticipated advances in the field of plasmonic DNA nanostructures, DNA nanostructures that respond to fluid flow and, eventually, modeling DNA nanostructures in the crowded environment of a biological cell.

**Figure 7:**
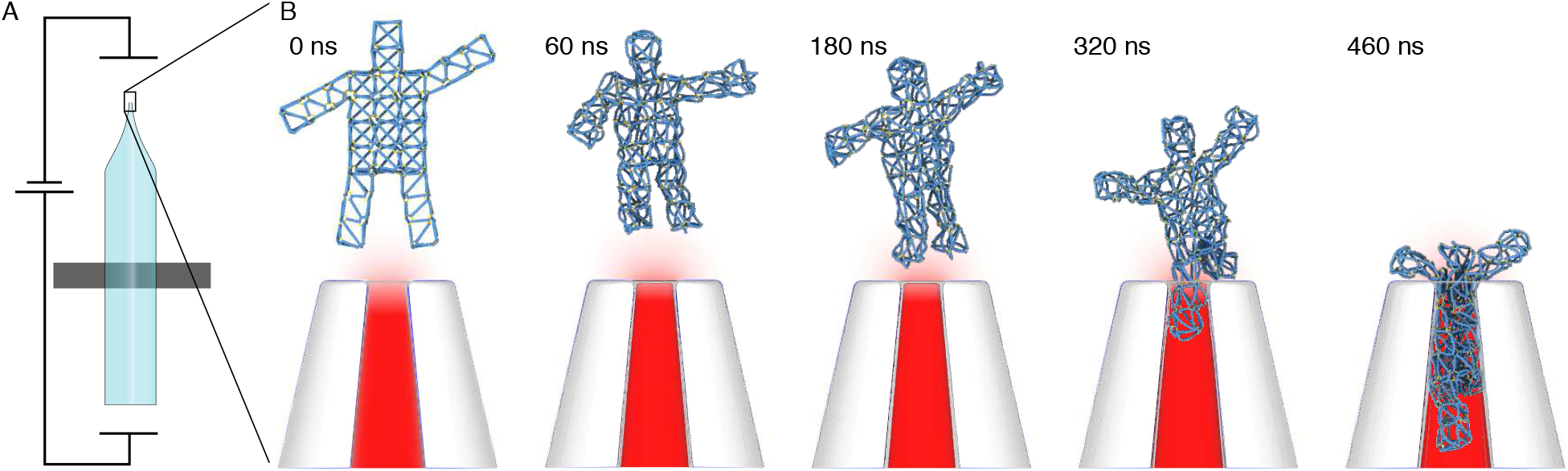
Coupling continuum models with `mrdna` simulations. (A) Initial configuration of a simulation system consisting of a vHelix-designed nanostructure^16^ near a glass nanopipette under a 300 mV applied bias. The electric potential distribution in and around the pipette was obtained using Comsol as previously reported^20^. The electric potential and a steric grid potential representing the presence of the nanopipette are incorporated in the `mrdna` model as external potentials. (B) `Mrdna` simulation of the DNA nanostructure capture by a nanopipette. From left to right, snapshots depict the configuration of the system during a 500-ns 5-bp/bead CG simulation.

## 4 CONCLUSION

We have presented an automated workflow for the prediction of the *in situ* structure and conformational dynamics of self-assembled DNA objects. Uniquely, our multi-resolution approach enables fast relaxation of an initial structural model through low-resolution CG simulations, progressive refinement of the structural models through finer-resolution CG simulations and, ultimately, provides a fully atomistic representation of the object’s *in situ* structure. In comparison to existing computational models of self-assembled DNA nanostructures, our multi-resolution approach is computationally much more efficient, providing relaxed, fully-atomistic models of typical 3D DNA origami nanostructures within 30 minutes or less. The quantitative accuracy of the resulting structural model allows the `mrdna` framework to be used routinely for DNA nanostructure design and prototyping, replacing labor- and time-consuming cryo-EM reconstruction. By taking into account non-trivial environmental conditions, such as variable ionic strength, applied electric field, non-homogeneous temperature and optical intensity, the framework enables computational study of, not only the structural, but also the functional aspects of the nanostructure designs.

The `mrdna` framework is distributed as an open source Python package^66^, enabling future customization of the algorithms by the end users. A command-line interface allows simulations to be performed and customized with a few keystrokes. The package provides additional capabilities to the users through simple Python scripting, such as the application of geometric transformations to parts of a DNA nanostructure, interfacing with continuum models through three-dimensional grid-specified potentials, and generating or modulating DNA nanostructures algorithmically. A step-by-step guide to using `mrdna` is available^67^.

Immediate future developments of the `mrdna` framework will focus on improving the description of ssDNA-dsDNA junctions, dsDNA nicks and end-stacking interactions. The physical realism of the model will be further enhanced by supplementing the `mrdna` framework with CG models that explicitly account for the sequence-dependence of DNA elasticity and the possibility of DNA hybridization and dehybridization reactions. Complementing development of ARBD, the framework will be extended to incorporate grid- and particle-based models of proteins and inorganic nano-structures. Finally, our multi-resolution simulation framework may be integrated with other DNA nanostructure design and visualization tools to provide near realtime feedback to the designer.

## Supporting information

Supporting Animation 1

Supporting Animation 2

Supporting Animation 3

Supporting Animation 4

Supporting Animation 5

Supporting Animation 6

Supporting Animation 7

Supporting Animation 8

Supporting Animation 9

Supporting Animation 10

Supporting Animation 11

Supporting Animation 12

## ACKNOWLEDGEMENTS

The authors gladly acknowledge supercomputer time provided through XSEDE Allocation Grant MCA05S028 and the Blue Waters supercomputer system (UIUC). The authors thank Shawn Douglas, Hendrik Dietz, Björn Högberg, Tim Liedl and many other practitioners of the structural DNA nanotechnology field for their valuable feedback on the pre-released version of the `mrdna` framework. A.A. thanks Dongran Han for sharing a cadnano design of the DNA flask. C.M. thanks Will Kaufhold for his help debugging the vHelix importer.

## FUNDING

This work was supported by the grants from the National Science Foundation (OAC-1740212 and DMR-1827346) and the National Institutes of Health (P41-GM104601).

## Conflict of interest statement

None declared.

