## Supplementary Information for "MrDNA: A multi-resolution model for predicting the structure and dynamics of nanoscale DNA objects"

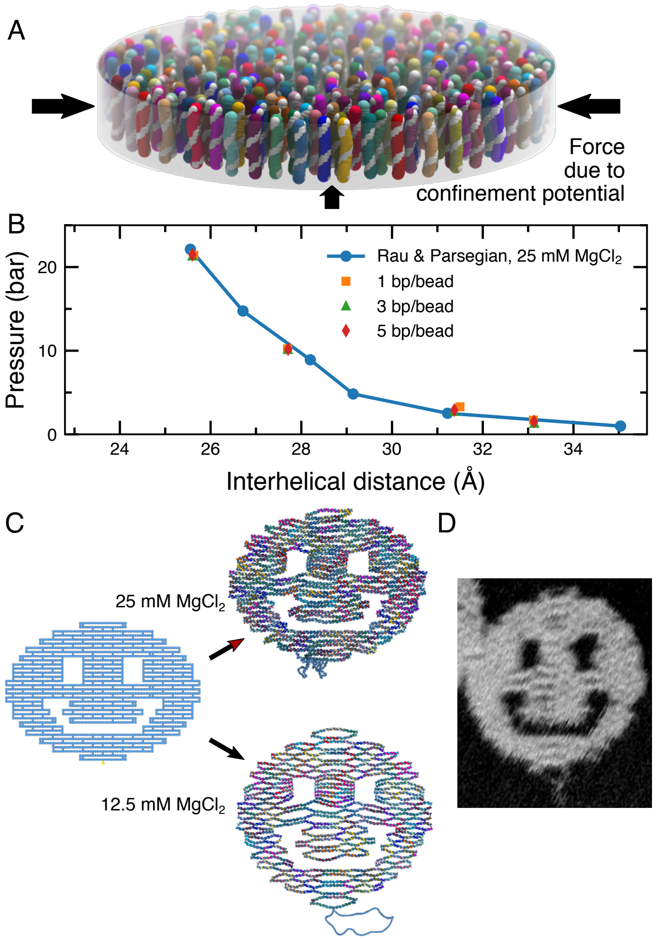


Figure S1: Calibration of non-bonded interactions. (A) Validation of the coarse-grained non-bonded potentials through simulations of a dsDNA array at several resolutions. The blue circles depict the experimentally derived pressure of a DNA array in 25 mM $MgCl_2$ solution as a function of the inter-DNA distance taken from Ref. 1. The colored symbols depict the pressure measured in `mrdna` simulations of 256 two-turn DNA helices at several resolutions. (B) The effect of ion concentration on the simulated structure of a two-dimensional DNA origami object. A cadnano file of the "smiley" object was produced according to the pattern described in the original manuscript.[2] The `mrdna` model of the smiley was relaxed from its initial configuration (left) using the default description of interactions in a 25 mM $MgCl_2$ solution (top) and using a Debye-Hückel correction to match the experimental conditions of 12.5 mM $MgCl_2$ (bottom). (C) Image of the smiley experimentally obtained using atomic force microscopy. Reproduced from Supplementary Figure S26 associated with Ref. 2.

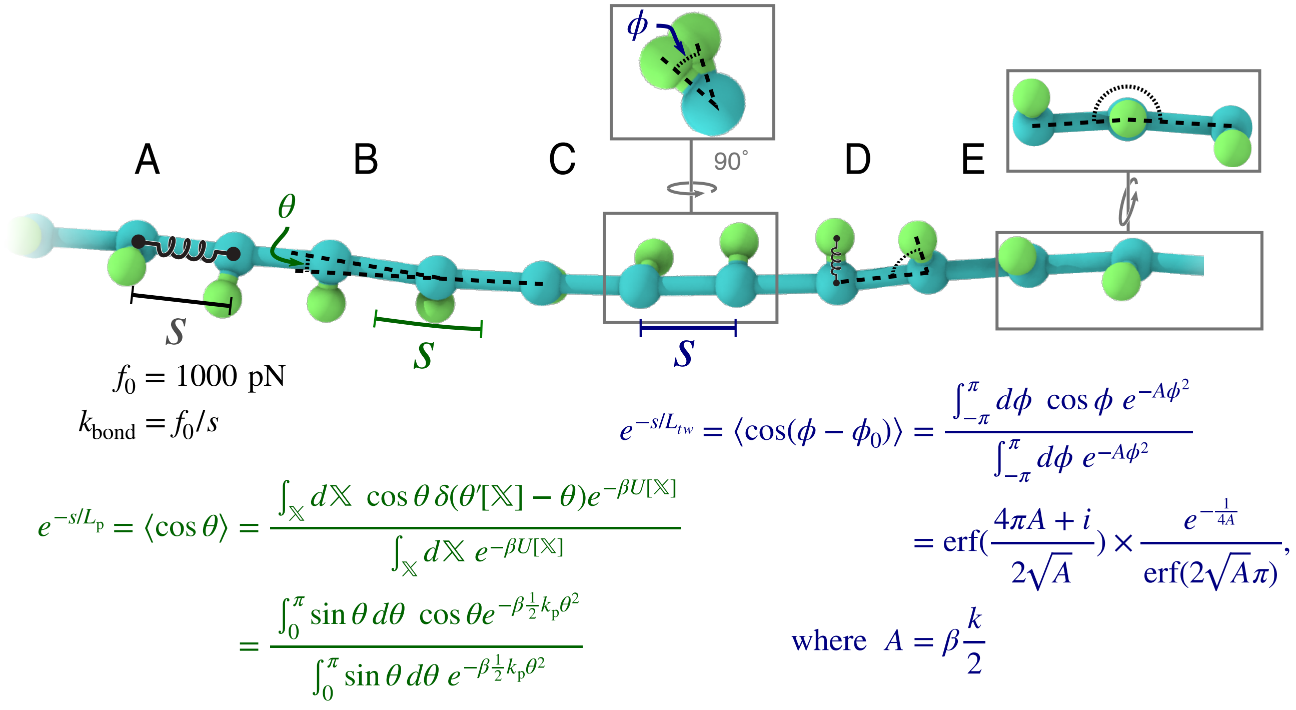


Figure S2: Schematic representation of bonded interactions in the `mrdna` model. (A) A harmonic spring connects adjacent beads with spring constant derived from the experimentally-determined elastic constant of dsDNA (1000 pN).[3] (B) A harmonic spring applied to the angle between the bonds formed by adjacent pairs of beads has its spring constant numerically determined from the experimentally-determined persistence length of dsDNA (50 nm). The remaining terms are applied when twist is locally represented. (C) A harmonic spring is applied to the dihedral angle formed by each orientation bead, its parent dsDNA bead, the adjacent dsDNA bead and its orientation bead to reproduce a twist persistence length within the range of experimentally-obtained values (90 nm).[4] (D) A harmonic bond associates each orientation bead with its dsDNA backbone bead (1.5 Å rest-length; $k_{\mathrm{spring}} = 30$ kcal/mol Å$^2$); the orientation bond is kept roughly normal to the local tangent of the DNA by a harmonic angle potential with 90° rest angle and $k_{\mathrm{spring}} = 0.25\ k_{\mathrm{L_p}}$; the backbone angle potential spring constant (schematically illustrated in panel B) is reduced by a factor of 0.75 to compensate for the added rigidity imparted by the orientation beads. (E) A harmonic potential applied to the improper dihedral angle formed by the bead below a dsDNA bead, the dsDNA bead, its orientation bead and the bead above the dsDNA bead provides additional stiffness in the direction normal to the bead axis to compensate for the loss of stiffness generated by reducing the spring constant of the angle potential (panel B) Black, green and blue equations correspond to the potentials described in (A–C), in that order.

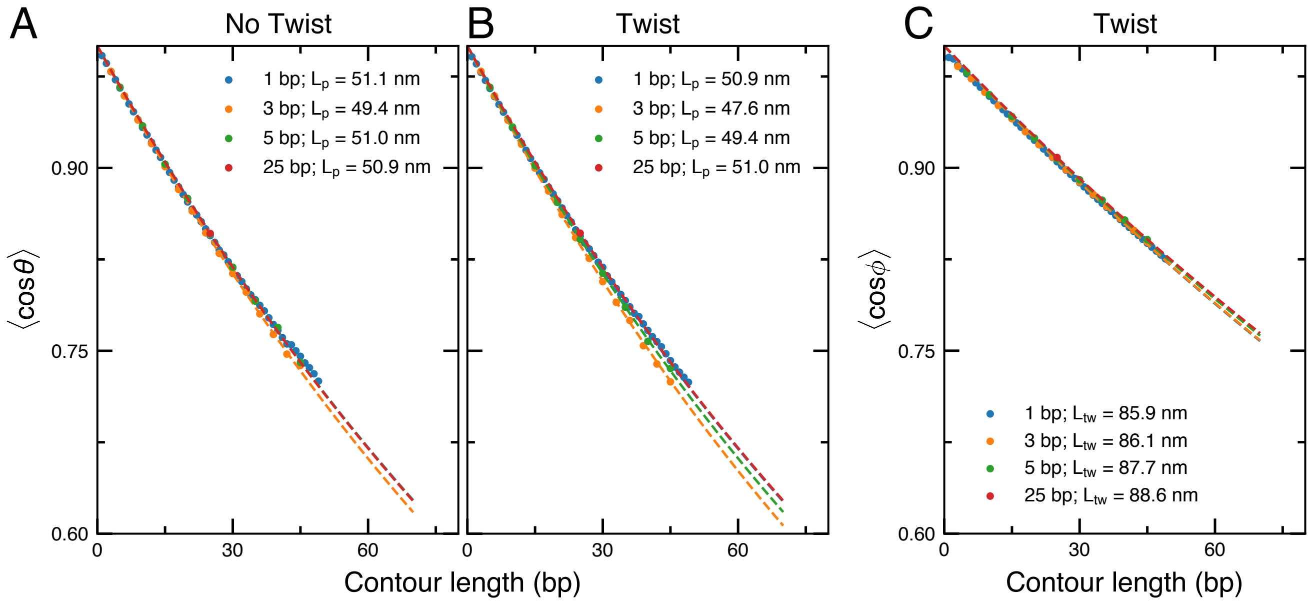


Figure S3: Polymer properties of the `mrdna` dsDNA model at different resolutions. (A,B) Contour-length dependence of the tangential correlations measured from `mrdna` simulations of a 300-bp dsDNA fragment at various resolutions lasting 500 million steps (40-fs timestep for 1-bp/dsDNA bead resolution; 150-fs otherwise), without (A) and with (B) a local representation of twist. Dashed lines depict exponentially decaying fits to the data, providing the persistence lengths of the DNA shown in the legend. (C) Contour-length dependence of the azimuthal correlation measured from the simulations described in panel B. Dashed lines depict exponentially decaying fits to the data, providing the twist persistence lengths of the DNA shown in the legend.

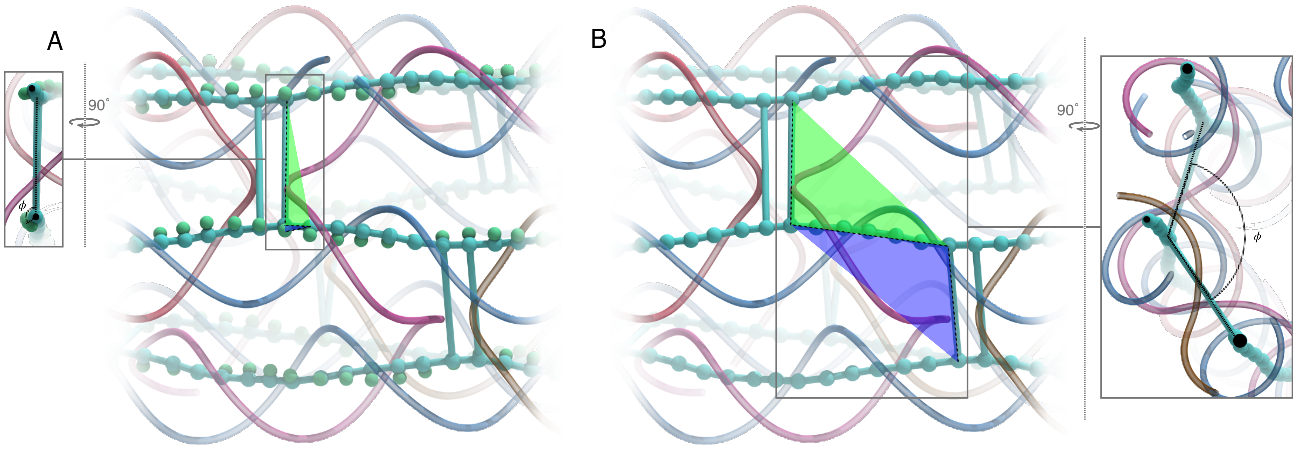


Figure S4: Twist in the `mrdna` dsDNA model around junctions. (A) Twist around dsDNA junctions when the model is constructed with a local representation of the DNA orientation. A harmonic potential is placed on the dihedral angle formed by two planes (green and blue in the figure) created by beads near the junction. The rest length is set to $\pm 120°$ with the sign depending on the strand. (B) Twist around dsDNA junctions when the model is constructed without a local representation of the DNA orientation. A harmonic potential is placed on the dihedral angle formed by two planes (green and blue in the figure) created by beads forming consecutive pairs of junctions. The rest length is set to $s \times 34.48°$, where $s$ is the contour length between the junctions in basepairs; the spring constant is derived from the twist persistence length of DNA.

### Supporting Animations

**Animation 1:** Movie illustrating the application of the `mrdna` framework for structure prediction of the pointer object[5] to obtain an all-atom model. The structure is depicted using the same representations as Fig. 2 of the main text.

**Animations 2-6:** Comparison of average simulated structures and cryo-EM reconstructed densities of the pointer object[5] (Animation 2); the v-brick structures without (Animation 3) and with the twist corrected (Animation 4);[6] and the rectangular (Animation 5) and triangular (Animation 6) prisms for hierarchical assembly.[6] Structures are depicted as in Fig. 3 of the main text.

**Animations 7,8:** Simulations of the flask object[7] designed by the Yan group before (Animation 7) and after (Animation 8) breaking the symmetry of the flask in a Python script. The structure

**Animations 9-11:** Depictions of structural fluctuations during 5-bp/bead resolution simulations of the caliper object from the Dietz group[8] (Animation 9); the slider object from the Castro group;[9] and the Bennett linkage object designed by the Castro group.[10]

**Animation 12:** Electrostatic capture of a wireframe mesh nanostructure[11] in a nanopipette with a 300-mV applied bias.
